# Modulating hinge flexibility in the APP transmembrane domain alters γ-secretase cleavage

**DOI:** 10.1101/375006

**Authors:** Alexander Götz, Nadine Mylonas, Philipp Högel, Mara Silber, Hannes Heinel, Simon Menig, Alexander Vogel, Hannes Feyrer, Daniel Huster, Burkhard Luy, Dieter Langosch, Christina Scharnagl, Claudia Muhle-Goll, Frits Kamp, Harald Steiner

## Abstract

Intramembrane cleavage of the β-amyloid precursor protein C99 substrate by γ-secretase is implicated in Alzheimer’s disease pathogenesis. Since conformational flexibility of a di-glycine hinge in the C99 transmembrane domain (TMD) might be critical for γ-secretase cleavage, we mutated one of the glycine residues, G38, to a helix-stabilizing leucine and to a helix-distorting proline. CD, NMR and hydrogen/deuterium exchange measurements as well as MD simulations showed that the mutations distinctly altered the intrinsic structural and dynamical properties of the TMD. However, although helix destabilization/unfolding was not observed at the initial ε-cleavage sites of C99, both mutants impaired γ-secretase cleavage and altered its cleavage specificity. Moreover, helix flexibility enabled by the di-glycine hinge translated to motions of other helix parts. Our data suggest that both local helix stabilization and destabilization in the di-glycine hinge may decrease the occurrence of enzyme-substrate complex conformations required for normal catalysis and that hinge mobility can be conducive for productive substrate-enzyme interactions.

## INSTRODUCTION

Proteolysis in the hydrophobic core of membranes is a fundamental cellular process that mediates critical signaling events as well as membrane protein turnover. Intramembrane proteases are found in all kingdoms of life and exist in several catalytic types (1). They are generally polytopic membrane proteins carrying their active sites residues in transmembrane helices (2). Apart from the fact that they cleave their substrates typically within the transmembrane domain (TMD), little is still known about the substrate determinants of intramembrane proteases as they do not, with few exceptions of some rhomboid proteases, recognize consensus sequences like common soluble proteases. Rather than sequence motifs, intrinsic instability and global flexibility of substrate TMD helices that could be induced by e.g. helix-destabilizing glycine residues in case of signal peptide peptidase (SPP) (3) or the related SPP-like (SPPL) protease SPPL2b (4), or a short helix-distorting asparagine-proline motif in case of site-2 protease (5, 6) are now increasingly discussed as a critical factor for substrate recognition and/or cleavage.

γ-Secretase is a pivotal intramembrane protease complex (7, 8) that cleaves more than hundred type I membrane protein substrates including signaling proteins essential for life such as Notch1 as well as the β-amyloid precursor protein (APP), which is central to the pathogenesis of Alzheimer’s disease (AD) (9, 10). It is widely believed that an aberrant generation and accumulation of amyloid β-peptide (Aβ) in the brain triggers the disease (11, 12). Aβ is a heterogeneous mixture of secreted, small peptides of 37-43 amino acids. Besides the major form Aβ_40_, the highly aggregation-prone longer forms Aβ_42_ and Aβ_43_ are pathogenic Aβ variants. Aβ species are generated by γ-secretase from an APP C-terminal fragment (C99) that originates from an initial APP cleavage of β-secretase (13). C99 is first endoproteolytically cleaved in its TMD by γ-secretase at the ε-sites close to the cytoplasmic TMD border and then processed stepwise by tripeptide-releasing carboxy-terminal trimming in two principal product lines thereby releasing the various Aβ species (14, 15). Mutations in the presenilins, the catalytic subunit of γ-secretase, are the major cause of familial forms of AD (FAD) and are associated with increased Aβ_42_ to total Aβ ratios (16). Rare mutations in the cleavage region of the C99 TMD also shift Aβ profiles and represent another cause of FAD (16, 17).

The molecular properties of substrates that are recognized by γ-secretase are still largely enigmatic (18). Established general substrate requirements are not only the presence of a short ectodomain (19, 20), which is typically generated by sheddases such as α- or β-secretase, but, equally important, also permissive transmembrane and intracellular substrate domains (18). Recent studies suggest that the recruitment of C99 to the active site occurs in a stepwise process involving prior binding to initial binding site(s) of the protease, so called exosites, in the nicastrin (NCT), PEN-2 and presenilin-1 (PS1) N-terminal fragment subunits of γ-secretase (21). Interactions with exosites may thus provide important checkpoints of the enzyme to distinguish substrates from nonsubstrates. Finally, at the active site, productive interactions of C99 with the S2’ subsite pocket of the enzyme are critical for substrate cleavage and Aβ product line selection (15).

Kinetic studies have shown that γ-secretase cleavage of C99 is a very slow process in the minute range, i.e. much slower than soluble proteases (22). Similarly low *k*_cat_ values as for C99 were determined for Notch1, which is unrelated in sequence to APP (19), and also for rhomboid proteases (23, 24). Low *k*_cat_ values together with a missing consensus sequence are indicative that conformational dynamics may govern substrate recognition (25). It is generally assumed that proteases cleave their substrates within extended sequences (i.e. β-strands) or loops (26). Although the cleavage sites can actually also reside in α-helices (27, 28), cleavable are mainly those helices that are intrinsically prone to unfolding or destabilization (29). Thus, the kinetics of the opening of the helix in cleavage domains of the TMD might be the key determinant for substrates. Additionally, local destabilization and the length of the membrane anchoring domains at the cytosolic juxtamembrane boundary, as well as β-sheet segments within the extracellular domain of C99 have been reported as important players for γ-secretase cleavage of C99 (30-32). This argues for conformational flexibility of the substrate being a key for productive interactions with the enzyme.

TMD helices are extremely stable due to their hydrogen bond network, which can be loosened by specific residues (33). For instance, the substitution of a single leucine by glycine in a membrane spanning poly-L helix facilitates helix bending, enhances local hydration, and triggers a redistribution of α-helical and 3_10_ helical H-bonds (34). The view that global TMD flexibility is required for efficient cleavage of C99 is further supported by biochemical analyses, which showed that mutations introduced at sites that are even farther from the cleavage region, can shift cleavage sites as well as cleavage efficiency (35-37). Furthermore, a detailed recent analysis showed that helix-instability of the C99 TMD caused by the introduction of helix-destabilizing di-glycine motifs in the domain near the ε-sites enhance the initial cleavage and subsequent carboxy-terminal trimming (38). Consistent with this view, biophysical studies indicate that substrate backbone dynamics could play a role in substrate selection (39). In particular the di-glycine hinge region in the C99 TMD (residues G37 and G38; Aβ numbering, see **Figure 1A**) has been suggested to provide the necessary flexibility for the interaction with the enzyme (25, 40-43). Thus, this G_37_G_38_ motif was shown to coordinate large scale bending movements of the C99 TMD (44) which might allow for a putative “swinging-in” of the C-terminal part of the helix to bring the scissile bonds in contact with the active site of γ-secretase (41, 45).

**Figure 1.**
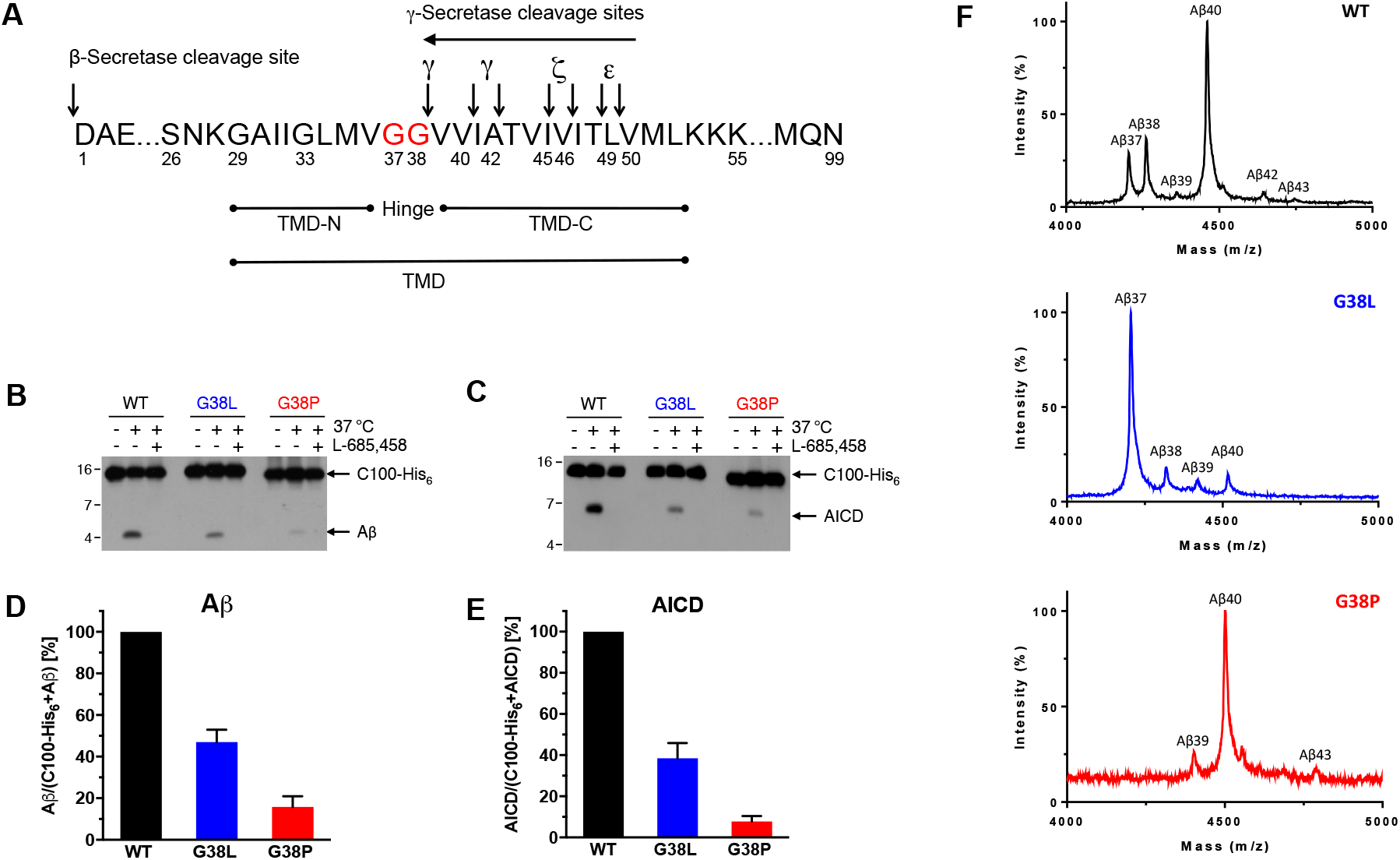
C99 G38P and G38L mutants distinctly alter γ-secretase cleavage and processivity. **(A)** Primary structure of C99 (Aß numbering) and its major γ-secretase cleavage sites. **(B)** Levels of Aß and **(C)** AICD were analyzed by immunoblotting after incubation of C100-His_6_ WT and mutant constructs with CHAPSO-solubilized HEK293 membrane fractions at 37 °C. As controls, samples were incubated at 4 °C or at 37 °C in the presence of the γ-secretase inhibitor L-685,458 (72). **(D)** Quantification of Aß and **(E)** AICD levels. Values are shown as a percentage of WT, which was set to 100%. Data are represented as mean ± SEM (n = 3, each n represents the mean of 3 technical replicates). **(F)** Representative MALDI-TOF spectra of the different Aß species generated for WT and the G38 mutants. The intensities of the highest Aß peaks were set to 100% in the spectra.

In order to evaluate the importance of hinge motions, we designed two artificial mutants of the C99 TMD with the aim to modulate the amount of hinge bending. A G38L mutation was intended to reduce bending by stabilizing the helix, due to the large hydrophobic side chain of leucine also allowing for van der Waals packing with the upstream L34 residue (46), while G38P should conversely increase bending considering the classical helix-breaking potential of proline in soluble proteins (47) and membrane proteins (48). In addition to γ-secretase cleavage assays of recombinant full-length C99 substrates, we assessed the effects introduced by the G38 mutations on the intrinsic structural and dynamical properties of peptides comprising the TMD of C99 and its C-terminal and N-terminal anchors (C99_26-55_, for sequences see **Table 1**) with several biophysical techniques and molecular dynamics simulations. C99_26-55_ peptides were reconstituted in model membranes to investigate the structure/dynamics of the C99 TMD in the hydrophobic environment. In addition, peptides were also studied in trifluoroethanol/water (TFE/H_2_O, 80/20 v/v), a classical solvent mixture to mimic the interior of proteins (49, 50), to adjust to the more hydrophilic environment inside γ-secretase (43, 44, 51-54).

**Table 1:**
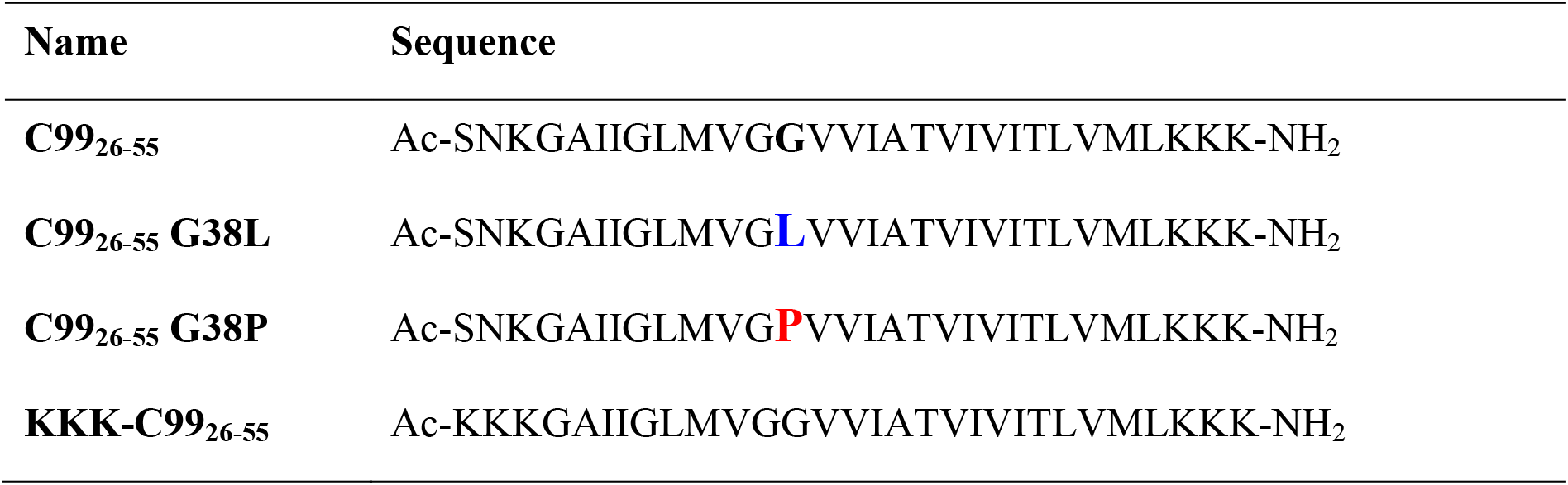
Sequences of investigated peptides.

We found that γ-secretase cleavage of both the G38L and particularly the G38P mutant of C99 was dramatically altered compared to WT. Thus, for both mutants total Aβ was reduced, and, remarkably, also the generation of the various Aβ species was distinctly altered for the mutants. The structural/dynamical studies of the C99 TMD peptides corroborated the expected “stiffening” and “loosening” effects of the G38L and G38P mutants, respectively, particularly in the TFE/H_2_O environment. Here, while the G38L mutant straightened, stiffened and increased the helicity of the C99 TMD, the G38P mutant indeed made the helix structurally less defined and more dynamic but introduced a structural kink located at the G_37_G_38_ sites. However, effects of the G38 mutations on the local dynamical properties of the C99 TMD were observed only in the vicinity of the G_37_G_38_ and did not extend to residues around the ε-sites at T48/L49, from which C99 cleavage by γ-secretase is initiated. Nevertheless, in both G38 mutants, the subtle changes in local H-bond flexibility in the vicinity of G_37_G_38_ introduced distinct large changes of intrinsic bending and twisting dynamics of the entire helix and distorted sampling of the orientation of the helical turn harboring the ε-sites. Altered global motions of the C99 TMD controlled by hinges around G_37_G_38_ may thus determine access to the active site enabling substrate-enzyme interactions required for catalysis.

## MATERIALS AND METHODS

### Materials

1-Palmitoyl-2-oleoyl-*sn*-glycero-3-phosphocholine (POPC) as well as *sn*-1 chain perdeuterated POPC-*d*_31_ were purchased from Avanti Polar Lipids. 1,1,1-3,3,3-Hexaflouroisopropanol (HFIP) and 2,2,2-trifluoroethanol (TFE) were purchased from Sigma Aldrich.

### Peptides

For all circular dichroism (CD), solution NMR, ssNMR and DHX experiments C99_26-55_, a 30 amino acid long peptide comprising residues 26 to 55 of C99 (C99 numbering, see **Figure 1A**), with N-terminal acetylation and C-terminal amidation was used. WT peptide and G38L and G38P mutants thereof (**Table 1**) were purchased from Peptide Specialty Laboratories GmbH, Heidelberg, Germany and from the Core Unit Peptid-Technologien, University of Leipzig, Germany. For ssNMR, A30, G33, L34, M35, V36, G37, A42, and V46 were labeled in the WT sequence with ^13^C and ^15^N. In the two mutant peptides, only A30, L34, G37, and V46 were labeled as a compromise between expensive labeling and highest information impact to be expected. For ETD measurements, in order to enhance fragmentation efficiency, we substituted the N-terminal SNK sequence of C99_26-55_ by KKK. In all cases purified peptides were to >90 % purity as judged by mass spectrometry (MS).

### γ-Secretase *in vitro* assay

C99-based WT and mutant substrates were expressed in *E. coli* as C100-His_6_ constructs (C99 fusion proteins containing an N-terminal methionine and a C-terminal His_6_-tag) (55) and purified by Ni-NTA affinity-chromatography. To analyze their cleavability by γ-secretase, 0.5 μM of the purified substrates were incubated overnight at 37 °C with CHAPSO-solubilized HEK293 membrane fractions containing γ-secretase as described (56). To control for specific cleavage by γ-secretase, substrates were incubated at 37 °C with 0.5 μM of the γ-secretase inhibitor L-685,458 (Merck Millipore) or incubated at 4 °C as a control. Generated Aβ and AICD was analyzed with immunoblotting using antibody 2D8 (57) and Penta-His, respectively, and quantified by measuring the chemiluminescence signal intensities with the LAS-4000 image reader (Fujifilm Life Science). Analysis of γ-secretase activity was repeated with three independent substrate purifications in three technical replicates for each of the constructs.

### Mass spectrometry analysis of Aβ species

Aβ species generated in the γ-secretase vitro assays were immunopreciptated with 4G8 antibody (Covance) and subjected to mass spectrometry analysis on a 4800 MALDI - TOF/TOF Analyzer (Applied Biosystems/MDS SCIEX) as described previously (58, 59).

### CD spectroscopy

C99_26-55_ WT, G38L and G38P mutant peptides were incorporated into large unilamellar vesicles (LUV) composed of POPC at a lipid/protein molar ratio of 30:1. First, 500 μg peptide and 3.72 mg POPC were co-mixed in 1 ml HFIP. After evaporation of the HFIP, the mixture was dissolved in 1 ml cyclohexane and lyophilized. The resulting fluffy powder was dissolved in 977 μl buffer (10 mM sodium phosphate, pH 7.4). After 10 freeze-thaw cycles, LUVs were prepared by extrusion using a 100-nm polycarbonate membrane and a LipofastTM extruder device (Armatis GmbH, Weinheim, Germany). CD spectra were recorded with a Jasco 810 spectropolarimeter. A cuvette with a 1 mm path length was filled with 200 μl of the LUV/C99_26-55_ preparation in which the final peptide concentration was 83 μM and the lipid concentration 2.5 mM. The UV absorbance at 210 nm of the WT peptide was used as a reference to normalize the final concentration of the reconstituted mutant peptides. Alternatively, the peptides (50 μM final concentration) were dissolved in TFE/H_2_O (80/20 v/v) and concentrations were determined based on the usage of the UV absorbance of the peptide bond at 205 nm with an extinction coefficient ε_205_ = 73.600 M^−1^cm^−1^. The latter was determined by calibration with a homologous peptide SNKWGAIIGLMVGGVVIATVIVITLVMLKKK whose concentration was determined using the ε_280_ = 5600 mol^−1^cm^−1^ of the additional tryptophan. Mean molar residue ellipticities ([Θ]) were calculated based on the peptide concentrations.

### Solution NMR

Dry C99_26-55_ WT (^15^N/^13^C-labeled at positions G29, G33, G37, G38, I41, V44, M51 and L52), G38L and G38P mutant peptides were dissolved in 500 μL 80% trifluoroethanol-d3 (TFE-d3) and 20% H_2_O respectively. pH was adjusted to 5.0 by adding the corresponding amount of NaOH. Peptide concentrations ranged between 50 to 500 μM. The NMR spectra of the peptides were obtained on a 600 MHz AVANCE III spectrometer (Bruker BioSpin, Rheinstetten, Germany) equipped with a TXI cryoprobe at a temperature of 300 K. To assign ^1^H and ^13^C-resonances of the peptides a set of two-dimensional spectra was recorded: ^1^H-^1^H-TOCSY with a mixing time of 60 ms, ^1^H-^1^H-NOESY with a mixing time of 200 ms, and ^1^H-^13^C-HSQC. Spectra were recorded with 24 scans and 1000 data points in the indirect dimension. The NMR spectra were analyzed using NMRViewJ (One Moon Scientific).

For hydrogen-deuterium (H/D) NMR-HDX exchange measurements dry peptides were dissolved in 80 % TFE-d3 and 20 % D_2_O. Measurements were done at at least three different pH-values to access all exchangeable protons, using the correlation of exchange rate and pH value. pH was adjusted using NaOD and DCl. Eleven TOCSY or ClipCOSY (60) spectra with an experimental time of 3 h 26 min each (mixing time 30 ms, 24 scans, 300 data points in the indirect dimension) were acquired sequentially. Additionally, eleven ^1^H-^15^N-HSQC spectra of the WT were recorded (2 scans, 128 points in the indirect dimension) in between.

The exchange of the first five to six and the last two residues was too fast to measure. M35 and A42 cross peak intensities were significantly lower than those of other amino acids. The H/D exchange rate constant (k_exp,HDX_) was obtained fitting the cross peak intensities over time to equation 1:

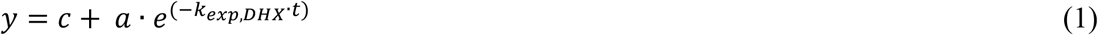

where t is time, a and c are constants. Rate constants were calculated for all three pH values and then scaled to pH 5.

### Solid-state NMR

Multilamellar vesicles were prepared by co-solubilizing POPC and the used C99_26-55_ peptide in HFIP at a 30:1 molar ratio. After evaporation of the solvent in a rotary evaporator, the sample film was dissolved by vortexing in cyclohexane. Subsequently, the samples were lyophilized to obtain a fluffy powder. The powder was hydrated with buffer (100 mM NaCl, 10 mM Hepes, pH 7.4) to achieve a hydration level of 50% (w/w) and homogenized by 10 freeze-thaw cycles combined with gentle centrifugation. Proper reconstitution of the C99_26-55_ WT and G38 mutant into the POPC membranes was confirmed by an analysis of the ^13^C C_α_ chemical shifts of A30.

^13^C magic-angle-spinning (MAS) NMR experiments were performed on a Bruker Avance III 600 MHz spectrometer (resonance frequency 600.1 MHz for ^1^H, 150.9 MHz for ^13^C) using 4 mm and 3.2 mm double-resonance MAS probes. The cross-polarization contact time was 700 μs, typical lengths of the 90° pulses were 4 μs for ^1^H, 5 μs for ^15^N and 4-5 μs for ^13^C. For heteronuclear two pulse phase modulation (TPPM) decoupling, a ^1^H radiofrequency field of 62.5 kHz was applied. ^13^C chemical shifts were referenced externally relative to tetramethylsilane. ^1^H-^13^C and ^1^H-^15^N dipolar couplings were measured by constant-time DIPSHIFT experiments using frequency-switched Lee Goldburg for homonuclear decoupling (80 kHz decoupling field) (61). The ^1^H-^13^C dipolar coupling was determined by simulating dipolar dephasing curves over one rotor period. These dipolar couplings were divided by the known rigid limit as reported previously (62). MAS experiments for the site-dependent order parameter were carried out at a MAS frequency of 3 kHz and a temperature of 30°C. DIPSHIFT ^1^H-^13^C order parameters were analyzed with a variant of the established GALA model (63) to evaluate RMSD values for combinations of tilt and azimuthal angles of the TMD helix, explained in detail in the Supporting Information.

### Mass spectrometric experiments of Deuterium/Hydrogen exchange (MS-DHX)

All mass spectrometric experiments were performed on a Synapt G2 HDMS (Waters Co., Milford, MA). A 100 μl Hamilton gas-tight syringe was used with a Harvard Apparatus 11plus, the flow rate was set to 5 μl/min. Spectra were acquired in a positive-ion mode with one scan for each second and 0.1 s interscan time.

Solutions of deuterated peptide (100 μM in 80 % (v/v) d1-trifluoroethanol (d1-TFE) in 2 mM ND_4_-acetate) were diluted 1:20 with protiated solvent (80% (v/v) TFE in 2 mM NH_4_-acetate, pH 5.0) to a final peptide concentration of 5 μM (at which the helices remain monomeric (43)), and incubated at a temperature of 20 °C in a thermal cycler (Eppendorf, Germany). At the used peptide concentration of 5 μM, the peptides remain monomeric (43). Incubation times were 0, 1, 2, 5, 10, 20, 30, 40, 50 min, and 1, 2, 3, 4, 6, 8, 12, 24, 48, 72 h. Exchange reactions were quenched by placing the samples on ice and adding 0.5% (v/v) formic acid, resulting in a pH ≈ 2.5. Mass/charge ratios were recorded and evaluated as previously described, (64, 65) including a correction for the dilution factor. For electron transfer dissociation (ETD) we preselected the 5+ charged peptides via MS/MS and used

1,4-dicyanobenzene as reagent. The fragmentation of peptides was performed as described. (64). Briefly, ETD MS/MS scans were accumulated over 10 min scan time, smoothed (Savitzky-Golay, 2 x 4 channels), and centered (80% centroid top, heights, 3 channels). MS-ETD-measurements were performed after 13 different incubation periods (from 1 min to 3 d) where exchange took place at pH 5.0. Shorter (0.1 min, 0.5 min) and longer (5 d, 7 d) incubation periods were simulated by lowering the pH to 4.0 or elevating pH to 6.45, respectively, using matched periods. The differences to pH 5.0 were considered when calculating the corresponding rate constants. We note that base-catalyzed exchange is responsible for at least 95 % of total deuteron exchange at pH 4.0 and above. The resulting ETD c and z fragment spectra were evaluated using a semi-automated procedure (MassMap_2017-11-16_LDK Software, MassMap GmbH &; Co. KG, Wolfratshausen) (39). The extent of hydrogen scrambling could not be calculated with the ammonia loss method due to the blocked N-termini. However, previous experiments with similar peptides showed scrambling to be negligible under our conditions (34). During all MS-DHX experiments, a gradual shift of monomodal shaped isotopic envelopes towards lower mass/charge values was observed. This is characteristic of EX2 kinetics with uncorrelated exchange of individual deuterons upon local unfolding (66, 67).

### Molecular Dynamics (MD) simulations

The sequences of the investigated model peptides are shown in **Table 1.** Because no experimental structures were available for the G38 mutants, we used a stochastic sampling protocol to generate a set of 78 initial start conformations (for details see (68)).

All-atom simulations in 80% TFE and 20% TIP3 (v/v) were set up as described previously (68). Each start conformation was simulated for 200 ns using (settings as described in (43)). Production runs were performed in a NPT ensemble (T = 293 K, p = 0.1 MPa) using NAMD 2.11 (69) and the CHARMM36 force field (70). The last 150 ns of each simulation were subjected to analysis, leading to an effective aggregated analysis time of 11.7 μs for each peptide. Frames were recorded every 10 ps.

For all-atom simulations in POPC bilayers, the 78 conformations were hierarchically clustered, the centroid of the cluster with the highest population was placed in a symmetric bilayer, consisting of 128 POPC lipids, using protocols as provided by CHARMM-GUI (71). Simulations of 2.5 μs (T = 303.15 K, p = 0.1 MPa) were performed, using NAMD 2.12 (69), the CHARMM36 force field (70) and settings as provided by CHARMM-GUI. Frames were recorded every 10 ps. Only the last 1.5 μs of the trajectory were subjected to analysis.

Analysis of the occupancies of the H-Bonds, tilt and azimuthal angles, as well as bending and twisting motions are explained in detail in the Supporting Information.

## RESULTS

### G38L and G38P mutations in the C99 TMD differently impair γ-secretase-cleavage

To examine whether and how leucine and proline mutations in the G_37_G_38_ hinge in the TMD of C99 impact on the cleavage by γ-secretase, G38L and G38P mutants of the C99-based recombinant substrate C100-His_6_ (55) were purified and used to assess their cleavability in an established *in vitro* assay (56). As expected, Aβ and AICD levels of the G38L mutant were reduced (~47% and ~38%, respectively) compared to WT (**Figures 1B, C, D** and **E**). Cleavage of the G38P mutant was even more reduced, leading to a residual Aβ and AICD production of ~16% and ~8%, respectively, compared to the WT. Thus, the hinge region is a critical part of the APP TMD, which, when mutated, can strongly influence substrate cleavability by γ-secretase. To also investigate the impact of the mutations on the carboxy-terminal trimming activity of γ-secretase, i.e. its processivity, we analyzed the finally released Aβ species with MALDI-TOF mass spectrometry (MS) (**Figures 1F**). Strikingly, while Aβ_40_ was as expected the predominant species for the cleavage of the WT substrate, Aβ37 was the major cleavage product for the G38L mutant. Thus, although the initial ε-cleavage was impaired for the G38L mutant, its processivity was enhanced. Aβ_40_ remained the major cleavage product for the G38P mutant, but in contrast to WT, no Aβ37 and Aβ38 species were produced. Additionally, also Aβ_43_ was detected for this mutant. Remarkably, for both mutants the unusual Aβ39 species was detected which was barely detected for WT. Thus, for both mutant substrates ε-cleavage and processivity by γ-secretase were markedly and distinctly affected.

### G38 hinge mutations cause structural changes of the C99 TMD helix

To understand why both hinge mutations impair γ-secretase cleavage, we next investigated how the G38 mutations affect structural and dynamical properties of the C99 TMD. To compare the effects of the G38L and G38P mutants on the helical conformation of the C99 TMD, WT and mutant peptides C99_26-55_ peptides (for sequences see Methods, **Table 1**), were incorporated into large unilamellar vesicles composed of POPC. As shown in **Figure 2A, CD** spectroscopy measurements demonstrated that the peptides contain a high content of α-helical conformation in the lipid bilayer. As expected, the helical conformation was stabilized for the G38L (indicated by the more negative ellipticity at 220 nm) and destabilized for the G38P mutant. Similar effects were found when the peptides were analyzed in TFE/H_2_O (80/20 v/v) (**Figure 2B**). In this solvent, ellipticity and the shape of the spectra was different compared to POPC, which was due to the different solvent properties, but the minima at 208 and 220 nm were nevertheless indicative of a high degree of helicity. The latter was corroborated by solution NMR (**Figure 2C**, see **Figure S1** for complete data set). Structural information was derived from NOE patterns and secondary chemical shifts (Δδ). Over the entire TMD sequence, C99_26-55_ WT showed cross peaks that are typical for an ideal α-helix (containing 3.6 residues per turn, **Figure S1**). Δδ(^13^C_α_) and Δδ(^1^H_α_) chemical shifts in particular, as well as Δδ(^13^Cβ) indicated a strong helicity for a C-terminal domain (TMD-C) ranging from V39 to L52 (**Figure 2C, Figure S1**). In contrast, the N-terminal domain (TMD-N), ranging from S26 up to V36, seemed to form a less stable helix. At positions G37 and G38 the helical pattern appeared to be disturbed which is obvious from the reduced values Δδ(^1^H_α_) at these residues. This observed pattern of TMD-N and TMD-C domains flanking the short G_37_G_38_ segment of lower stability is consistent with previous results (43, 44, 53, 73).

**Figure 2.**
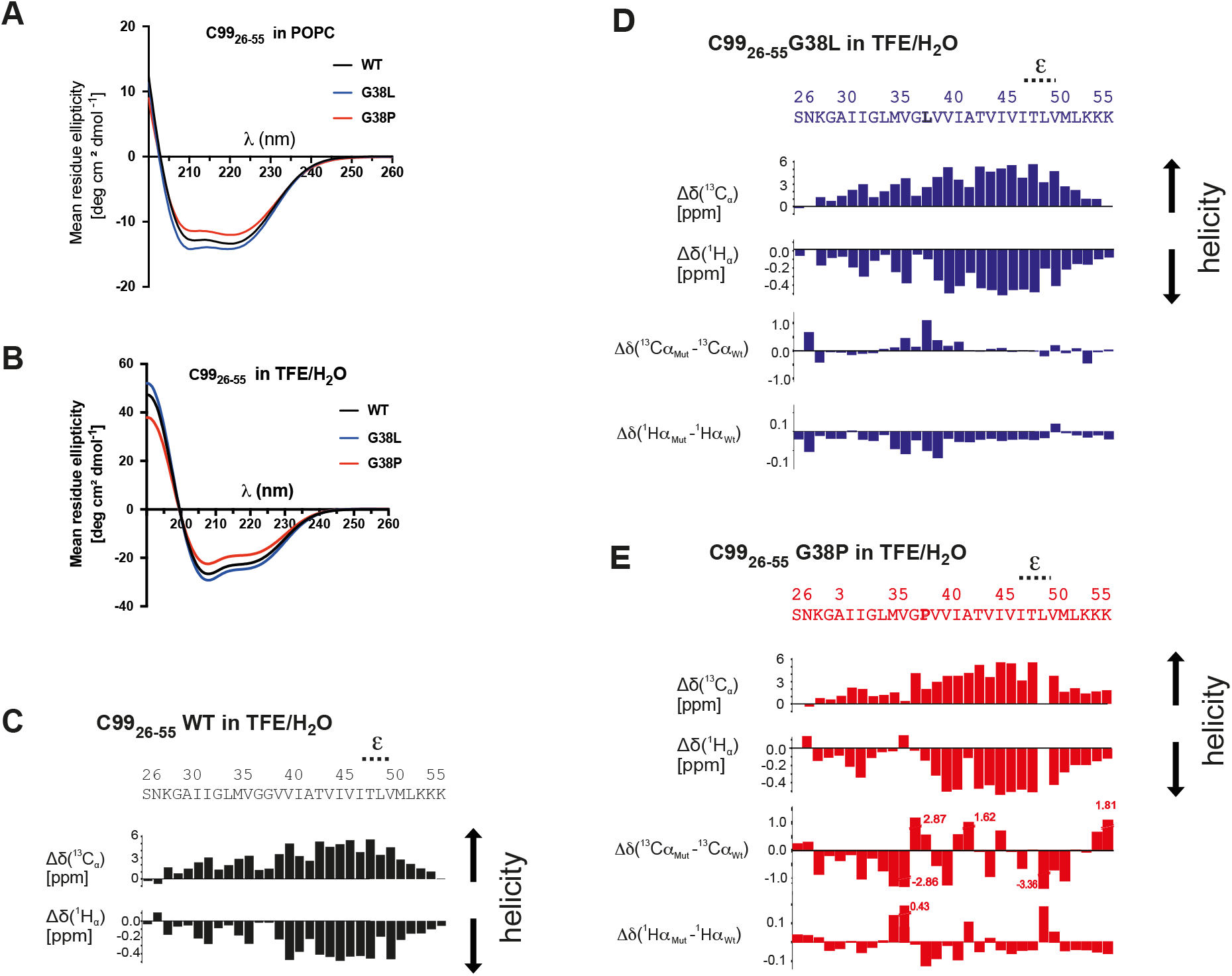
Helicity of C99_26-55_ TMD peptides is increased by the G38L and distorted by the G38P mutation. **(A)** CD spectra of C99_26-55_ WT, G38L and G38P mutant peptides reconstituted in POPC model membranes and **(B)** dissolved in TFE/H_2_O. **(C)** Chemical shift indices (Δδ) for ^13^C_α_ and ^1^H_α_ atoms of each residue of C99_26-55_ WT obtained from solution NMR in TFE/H_2_O. **(D)** and **(E)** Results of solution NMR measurements as in (C) of the G38L and G38P mutants, respectively, where also the differences between Δδ values of mutants and WT are depicted.

With regard to the G38 mutants, no major changes in the NOE patterns and secondary chemical shifts were observed. However, a detailed view showed small differences in secondary chemical shifts both for G38L and G38P. G38L appeared slightly stabilized compared to the WT. Chemical shift changes were restricted to the two helical turns around L38 (M35 - I41) and the immediate termini (**Figure 2D**). For G38P the high number of differences in both Δδ(^13^C_α_) and Δδ(^1^H_α_) shifts compared to those of the WT values (**Figure 2E**) indicated changes in structure or stability induced by the mutation that, however, were too subtle to also result in altered NOE patterns. This stems from the fact that NOEs for dynamic helices are dominated by the most stable conformation, whereas the highly sensitive chemical shifts are affected even by minuscule changes. Particularly the helicity of the N-terminal part up to V36 had decreased according to the chemical shift pattern, as predicted, but also the remaining C-terminal part showed significant and irregular deviations compared to the WT. Concomitantly, for G38P we observed an overall increase of H_N_ resonance line widths, which is indicative of an increased global conformational exchange (**Figure S2**). Here, also several minor alternative conformations were visible, that resulted in more than one H_N_/H_α_ resonance for many residues (up to four for some residues) (**Figure S2**). Taken together, consistent with the CD data, the solution NMR data show that the C99_26-55_ peptide has a high propensity to form a helical structure, which is only slightly reduced in the G_3_7G_38_ hinge region. Interestingly, the TMD-C, i.e. the region in which the cleavages by γ-secretase occur, has a much stronger helicity compared to the TMD-N. The G38L mutation caused a stabilization of the helix around the G_37_G_38_ hinge, albeit small, whereas the G38P mutation disturbed the helix both in its TMD-N and TMD-C parts.

### G38 mutant helices display altered hydrogen-bond stability around the G_37_G_38_ hinge

We next assessed the conformational flexibility of the C99_26-55_ WT and mutant helices, as expressed by intrahelical amide H-bond stabilities. To this end, we performed backbone amide deuterium-to-hydrogen (D/H) exchange experiments in TFE/H_2_O using MS (MS-DHX) as well as hydrogen-to-deuterium (H/D) exchange using NMR (NMR-HDX). Determining amide exchange in POPC membranes was not feasible, as the bilayer effectively shields central parts of the TMD helix (64, 65) so that backbone amides engaged in H-bonds do not exchange. Generally, although exchange rate constants also depend on the local concentration of hydroxyl ions (i.e. the exchange catalyst) and are influenced by side-chain chemistry (66), the reduced stability of backbone amide H-bonds associated with more flexible helices results in faster amide exchange. **Figure 3A** shows the result of MS-DHX experiments of >98% deuterated C99_26-55_ WT and mutant peptides (5 μM) in TFE/H_2_O. Consistent with previous results (43, 44, 52), overall D/H exchange was characterized by rapid deuteron exchange within minutes followed by a relatively fast exchange over 60 min (**Figure 3A**, inset) and a subsequent very slow process; near complete exchange was seen after three days. Relative to WT, the G38L mutation slowed hydrogen exchange. G38P resulted in a slightly faster exchange. In addition, we detected a reproducible decrease of a further 0.5 D after a few minutes of exchange (**Figure 3A**, inset) suggesting acceleration of exchange by the G38P mutation.

**Figure 3.**
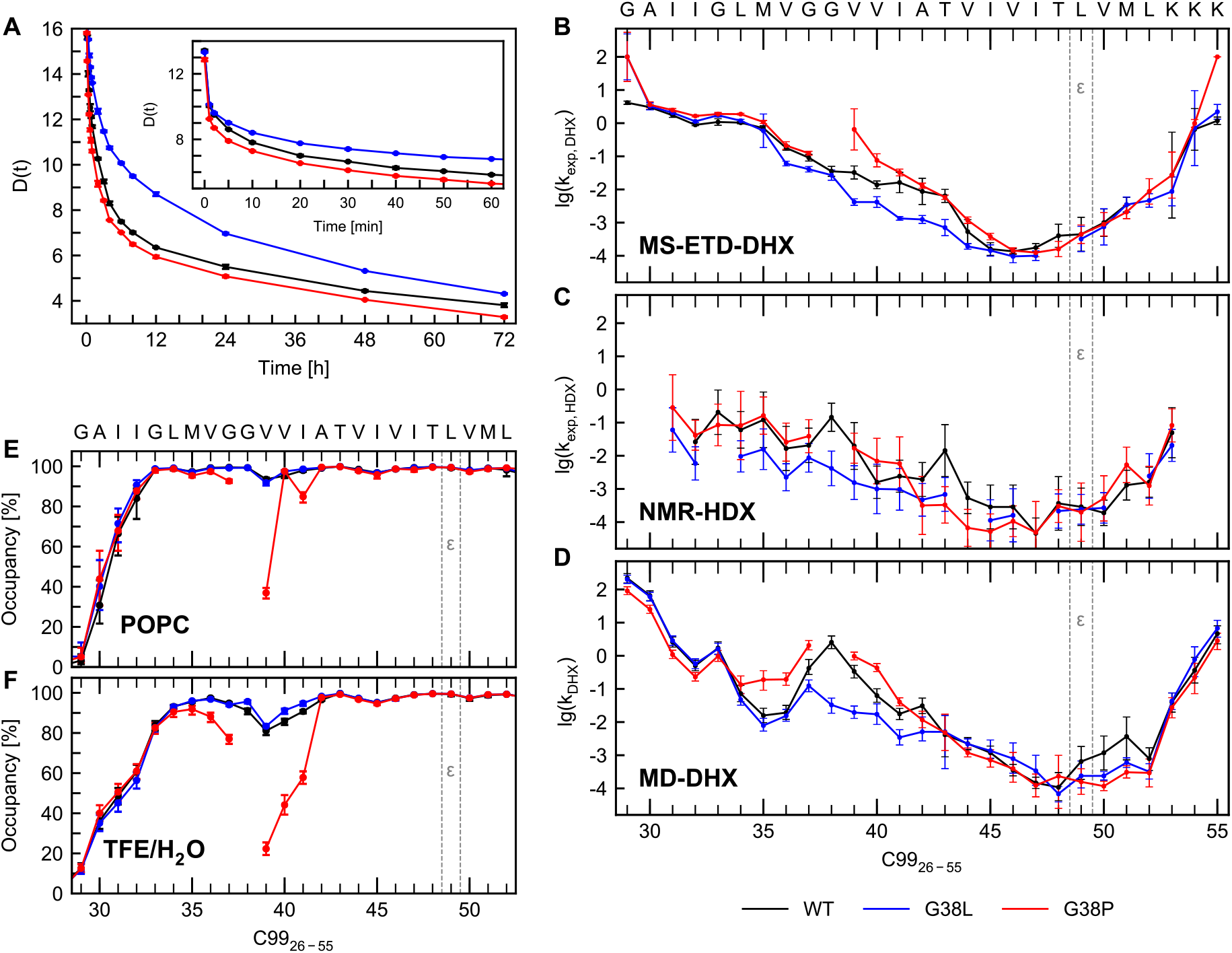
Deuterium-hydrogen exchange rates along the TMD of C99_26-55_ reveal an impact of the G38 mutations on H-bond stability around the mutation sites, but not at the ε-sites. **(A)** Overall DHX kinetics of C99_26-55_ WT, G38L and G38P mutant peptides measured with MS-DHX. Complete deuteration was followed by back-exchange in TFE/H_2_O, pH 5.0, T = 20 °C. Exchange kinetics during the first 60 min (inset) and 72 h were measured (n = 3, error bars showing SD are smaller than the size of the symbols). Note that a part of the lower deuterium content in G38P resulted from the lack of one amide deuteron at the cyclic side chain of proline. **(B)** Site specific D/H exchange rate c367 cl:1onstants (k_exp_,_DHX_ [min^−1^]) of C99_26-55_ WT, G38L and G38P mutants dissolved in TFE/H_2_O as determined by MS-ETD (error bars show 95% CI). **(C)** Site specific H/D exchange rate constants (k_exp,HDX_ [min^−1^]) determined by NMR. (n = 3, error bars show SD). **(D)** Site-specific k_DHX_ [min^−1^] computed from MD simulations (error bars show 95% CI). **(E)** Backbone H-bond occupancy of the individual residues of C99_26-55_ WT and its G38L and G38P mutants in POPC and **(F)** in TFE/H_2_O, calculated by MD simulations. Note that G38P cannot not form H-bonds at residue 38 due to the chemical nature of proline.

To obtain insight into local amide H-bond strength, we next measured residue-specific amide D/H exchange rate constants (k_exp,DHX_) using electron transfer dissociation (ETD) of our TMD peptides in the gas-phase after various periods of exchange (MS-ETD-DHX) (34, 39). In order to enhance fragmentation efficiency, we substituted the N-terminal SNK sequence of the C99_26-55_ TMD by KKK. As shown in **Figure 3B**, for all three peptides, D/H-exchange occurred within minutes for residues up to M35 within TMD-N and at the C-terminal KKK residues (68). The rate constants gradually decreased by up to two orders of magnitude in the region harboring the G_37_G_38_ motif. Interestingly, compared to WT, both G38 mutants perturbed exchange downstream of the mutation site in the region around the γ-40 cleavage site. While the G38L mutation decreased k_exp,DHX_ significantly between V39 and T43, G38P increased k_exp,DHX_ mainly for V39 and V40. Very slow exchange was observed in TMD-C, containing the ε-cleavage sites, which was not affected by the G38 mutants. Additionally, we measured H/D exchange by NMR spectroscopy. The shape of the NMR-HDX profile (**Figure 3C**) roughly matched the MS-ETD-DHX profile in that exchange within the TMD-N was faster than within TMD-C (**Figure 3B**). Further, NMR confirmed locally reduced exchange rates for the G38L mutant and locally enhanced rates for G38P, although the experimental errors prevented clear assignments of the differences to individual residues. As for MS-ETD-DHX, the G38 mutants did not affect the NMR-HDX in the vicinity of the ε-cleavage sites. The generally lower H/D rate constants, relative to the respective D/H values, are ascribed to the intrinsically slower chemical H/D exchange as compared to D/H exchange (68, 74).

Furthermore, DHX rate constants reconstructed by MD simulations from the fraction of open H-bonds and the local water concentration could reproduce the overall MS-DHX kinetics well (0.400 ≤ *χ*^2^ ≤ 1.493, **Figure S3**). In accordance with the ETD-derived rate profile, the calculated site-specific k_DHX_ exchange rate constants (**Figure 3D**) revealed fast exchange at both termini and very slow exchange in TMD-C. Additionally, the slow exchange at the ε-sites, without significant differences between WT and the G38 mutants was confirmed. Taken together, for all peptides, local amide exchange rates determined by three different techniques consistently reported perturbed backbone stability in the helix-turn downstream to the G38 mutation site in the γ-cleavage site region, but no alterations in helix stability around the ε-sites.

To gain further insight in the distribution of flexibility along the C99_26-55_ peptides we focused on the site-specific population of α-H-bonds (NH(i)…O(i-4)) and 3_10_-H-bonds (NH(i)…O(i-3)). Membrane proteins can show backbone H-bond shifting that does not induce permanent conversion to alternative helical forms (e.g. 3_10_-helix, which has only 3 residues per turn) but rather confers flexibility to the TMD helices and provides pathways for conformational changes like helix bending and twisting that accompany functional cycles (33). Since switching between α- and 3_10_-H-bonds has been detected previously for C99_28-55_ (43, 44, 53) as well as for other TMDs (32, 34), we calculated both α- and 3_10_-H-bond occupancies for each residue of C99_26-55_ in POPC and TFE/H_2_O, respectively, from the MD simulations (**Figure S4**). For the WT and G38L peptides in POPC a 10% lower occupancy of α-H-bonds emanating from backbone amides of V39 and V40 was largely compensated through the formation of 3_10_-H-bonds (**Figure S4**). In TFE/H_2_O a larger stretch of H-bonds spanning from the G33 carbonyl-oxygen to the amide-hydrogen at I41 was destabilized (**Figure S4**). Here, a maximal drop in α-helicity by 40% was only partially compensated through the formation of 3_10_-H-bonds indicating enhanced conformational variability. We also calculated the combined occupancies from **Figure S4** where an amide is regarded as protected from exchange if either the α- or the 3_10_-H-bond is formed. As shown in **Figures 3E** and **F**, the resulting occupancy loss for these H-bonds correlated with flexibility at the G_37_G_38_-hinge where H-bonds on the opposite face of the hinge have to stretch in order to allow for bending. Thus, it was found that only around the G3_7_G_38_ the H-bonds were distorted.

With regard to the ε-sites, in the TMD-C of all C99_26-55_ peptides both in POPC and TFE/H_2_O, we found a 5-10% population of 3_10_-H-bonds around the amides of T43/V44 and T48/L49 (**Figure S4**). However, neither shifting between α- and 3_10_-H-bonds nor helix distortions involving the carbonyl-oxygen at the ε-sites or other signs of helix distortions could be detected for the G38 mutants (**Figures S4** and **3E, F**). Remarkably, the occupancies of 3_10_-H-bonds did not change when changing solvent from POPC to the hydrophilic environment in the TFE/water solution. This demonstrates that the ε-sites are in a rather stable helical conformation, regardless of the solvent, which is not perturbed by the G38 mutations.

### G38 mutants alter the spatial orientation of the ε-cleavage site region

In addition to local variations of the structure and stability of the C99 TMD investigated above, it is possible that also global orientation of the TMD helix in the bilayer plays an important role in substrate recognition and cleavage. TMD helices usually tilt in order to compensate hydrophobic mismatching between the length of the hydrophobic domain of the TMD and the hydrophobic thickness of the lipid bilayer (75, 76). Tilt angles fluctuate with time. Additionally, azimuthal rotations of the tilted helix along its axis are not randomly distributed but reflect preferential side chain interactions with the individual components of the phospholipid bilayer (76-78). To explore a potential influence of the G38 mutations on these properties, we investigated the distribution of tilt (τ) and concomitant azimuthal rotation (ρ) angles (**Figure S5A**) of C99_26-55_ embedded in a POPC bilayer by solid-state NMR (ssNMR) and MD simulations. The C_α_-H_α_ order parameters of residues A30, G33, L34, M35, V36, G37, A42 and V46 of C99_26-55_, chosen to represent the helical wheel with C_α_-H_α_ bond vectors pointing in many different directions, were derived from DIPSHIFT experiments, which also confirmed the proper reconstitution of the peptides in the POPC bilayer (**Figure S6**). In order to estimate τ and ρ of the TMD helix in C99_26-55_, the GALA model (63) was used (Supporting Information).

In **Figure 4A** the normalized inverse of the root-mean-square deviation (RMSD_Norm_) between data and model is shown as function of tilt and azimuthal rotation angles. For all three peptides relatively broad τ, ρ landscapes with several possible orientations were found. Reliable helix orientations were found for all three peptides comprising helix tilt angles τ on the order of 30° or below, although the G38P landscape deviated from the similar landscapes of WT and the G38L mutant. An averagely small tilt angle for WT C99 TMD was also found by others (79), although in the latter study a very heterogeneous picture with different orientation and dynamics of several helix parts was drawn. As shown in **Figure 4B**, the calculated probability distributions of (τ,ρ) combinations from MD simulations were in agreement with the ssNMR observations. Average tilt angles were in the order of 30° or below (WT: 23.1° with 95% CI [20.2°, 26.2°] in agreement with previous results (40), G38L: 21.8° ([18.7°, 25.2°]) and G38P 25.9° ([22.2°, 29.9°]). A precise average ρ angle could not be calculated from the order parameters obtained from ssNMR nor obtained from the MD simulation, but only a range of possible orientations, which is indicative of high TMD helix dynamics in liquid-crystalline membranes.

**Figure 4.**
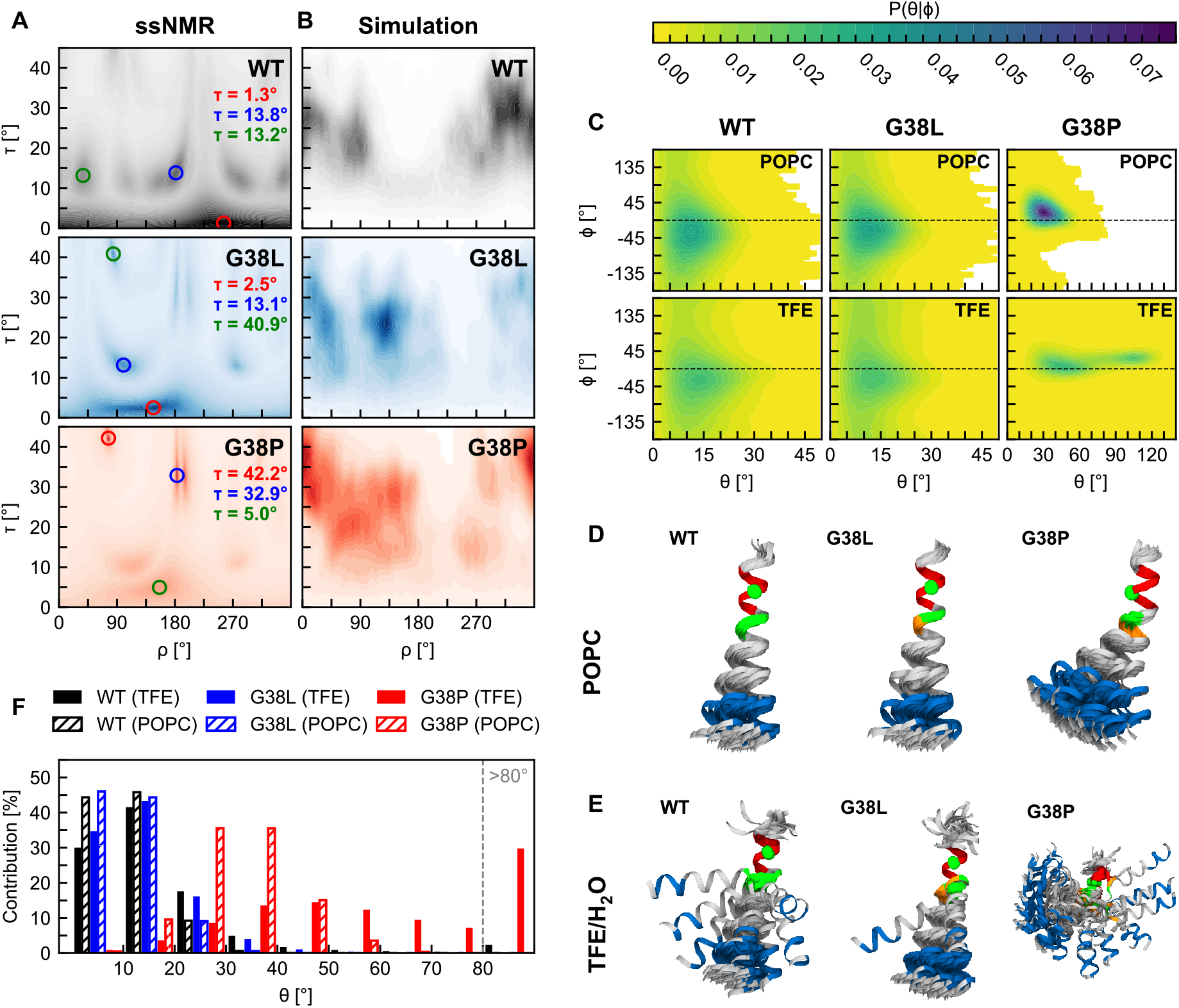
G38 mutations of the C99_26-55_ peptide do not significantly alter its membrane orientation but change orientation of the ε-sites region. **(A)** Heat maps of tilt (τ) and azimuthal rotation (ρ) angle combinations of C99_26-55_ WT, G38L and G38P mutant peptides in a POPC bilayer as determined by ssNMR. The colours represent the RMSD_Norm_ of the given (τ, ρ) pair. Maxima (dark areas) represent possible orientations. The circles represent the likeliest (red), second likeliest (blue) and third likeliest (green) solutions. **(B)** Probability distributions P(τ,ρ) of τ and ρ angle combinations of C99_26-55_ WT, G38L and G38P mutants in a POPC bilayer calculated from MD simulations. Dark areas represent high probabilities. **(C)** Probability distributions of bending (θ) and swivel (φ) angle combinations characterizing the orientation of ε-sites in C99_26-55_ WT, G38L and G38P mutants in POPC and in TFE/H_2_O, calculated from MD simulations. (**D, E**) Representative conformations for WT and G38 mutants in **(D)** POPC and **(E)** TFE/H_2_O determined by K-means clustering of (θ,φ) combinations in cos/sin space. Domains colored in blue indicate the TMD-C segment I47-M51 carrying the ε-sites. Domains colored in red represent the TMD-N segment I31-M35, which was also used to overlay the structures. The G_37_G_38_ motif is colored in green. For the G38 mutants the L and P residues are depicted in orange. Green spheres represent the C_α_ atom of G33 used as reference for the determination of swivel angles. **(F)** Distribution of conformations according to their bending angles θ. The last class summarizes all conformations with θ > 80°.

With respect to the reduced cleavability of the G38 mutants, the orientation of the ε-sites, at which the substrate is cleaved first, is of special interest. We thus investigated the orientation of the helical turn carrying the ε-sites (domain B: I47-M51, colored blue in **Figures 4D** and **E**) with respect to the orientation of the helical turn in TMD-N around G33 (domain A: I31-M35, colored red in **Figures 4D** and **E**) by a pair of angles. The bending angle (θ) is calculated as the angle between the axes through the two segments. The swivel angle (φ) is defined by the horizontal rotation of the domain B relative to domain A (**Figure S5B**). Positive φ-angles represent counter-clockwise rotation. We analyzed the bend and swivel conformational sampling from the MD simulations for the C99_26-55_ WT and G38 mutant peptides in POPC and TFE/H_2_O and report probability distributions of (θ,φ) combinations in **Figure 4C** as well as average bend/swivel geometries in **Table S1**. Representative conformations are exemplified in **Figures 4D** and **4E**.

Generally, we observed an asymmetry of the bend-swivel behavior of the ε-site orientations in all peptides with respect to both, structure and dynamics. In POPC, the ε-sites-containing domain B of the WT and G38L peptides exhibited bending rarely exceeding 30° with a mean value of ~12°. For the G38P mutant, an increased population of conformations with θ even larger than 40°, lack of conformations with θ<15° (**Figure 4F**), and an average bending of ~32° reveal a persistent reorientation of the ε-sites. However, the range of bending angles sampled around the mean value is ~30° for WT as well as for mutant peptides. Examination of the swivel angles reveals anisotropic bending, where the preference for particular regions of the swivel space is determined by residue 38. Both G38 mutations impart a counterclockwise shift of the sampled swivel angle space. As a consequence, the mean φ angle shifts by 10° for the G38L mutant and by 40° for the G38P mutant with respect to the WT (**Table S1**). Compared to WT and G38L peptides, which sample a swivel angle range of ~100°, the G38P mutation favors a much narrower range of swivel angles (~60°). Changing from the POPC membrane to TFE/H_2_O does not shift preference of the peptides for the particular regions in the swivel space (**Figure 4C**). However, we generally note an increase in the fraction of conformations with larger bending angles (**Figure 4F**). In the case of the G38P peptide, we even notice excursions to a population with large bending angles θ>80° (**Figure 4E, Table S1**).

Taken together, the comparison of the bend/swivel behavior of ε-site orientations revealed an asymmetry in helical conformations. The G38P mutation alters both bend and swivel angles, i.e. vertical and horizontal position of the ε-cleavage site region. In contrast, the G38L mutation mainly impacts the swivel angles and thus misdirects the ε-cleavage region horizontally. Additionally, the broader distributions of tilt and azimuthal rotation angles observed for G38P and the slightly narrower distributions for G38L compared to WT, as found in the MD simulations and also experimentally, might not only indicate differences between these mutants in their intrinsic membrane orientation but could also reflect the increased bending of G38P.

### G38 mutations relocate hinge sites and alter extent of hinge bending and twisting

The results discussed so far revealed that the impact of the G_37_G_38_-hinge mutations on H-bond stability is confined to a small number of residues in the hinge region. Although H-bonding around the ε-sites was not altered, we noticed that sampling of ε-site orientations is distorted in the mutants. Generally, TMD helices bend and twist by changing the direction of the helix axis or the helical pitch around various flexible sites (80-83). Since all these helix distortions may contribute to the variability of the orientation of the ε-cleavage site region (**Figure 4C**), we next analyzed from MD simulations the six fundamental types of helix motions including bending or twisting around a single hinge (referred to as types B and T) and combined bending and/or twisting around a pair of hinges (referred to as types BB, BT, TB and TT) (34, 53). These six types of subdomain motions are exemplified in **Figure 5A** for C99_2_6-55 WT.

In order to understand the impact of the G38 mutations on the variability of the orientations of the ε-cleavage site region, we next investigated the subdomain motions in WT and mutant peptides. Hinge sites are detected as flexible joints, able to coordinate the motion of more rigid flanking segments (34, 53, 68, 84). The contribution of each type of subdomain motion is depicted in **Figures 5B** and **C**. More than 90% of the sampled conformations deviated from a straight helix. In the POPC bilayer, bending and twisting around a single hinge (types B+T) contributed ~55% to overall backbone flexibility, motions around a pair of hinges (types BB, BT, TB and TT) contributed ~35%. Note that the same residues provide bending as well as twisting flexibility. The loss of packing constraints from lipids as well as enhanced H-bond flexibility in TFE/H_2_O (**Figures 3E, F**) correlated with favored helix bending (types B and BB) in WT and G38L. Remarkably, for G38P both single bending (type B) as well as twisting (type T) around the G_37_G_38_ motif were enhanced on the expense of all other hinge motions.

**Figure 5.**
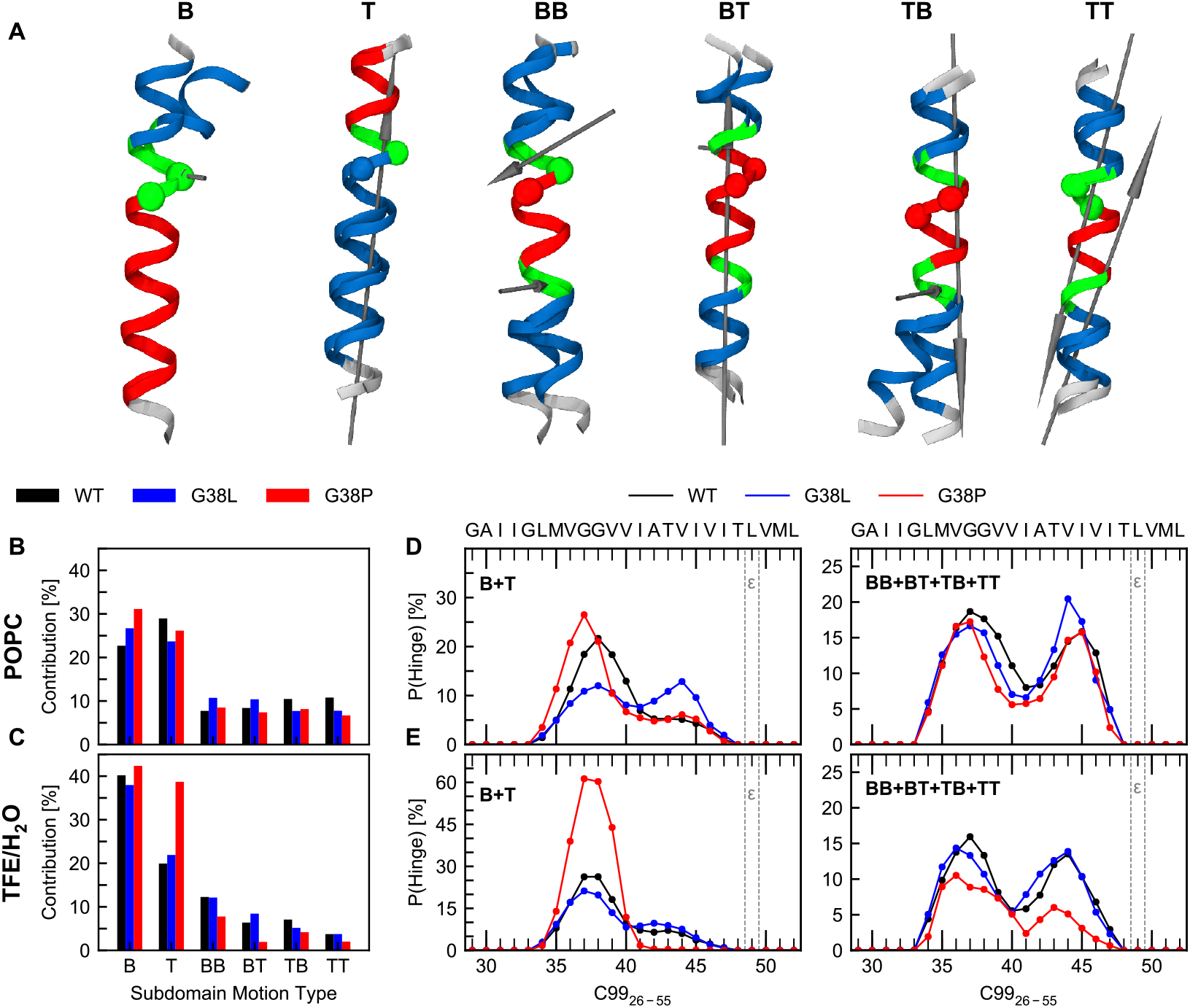
G38 mutations alter global bending and twisting motions. **(A)** The fundamental motions of helices exemplified for the C99_26-55_ WT peptide. Motion types are bending (B) and twisting (T) coordinated by a single hinge as well as combinations of bending and twisting (types BB, BT, TB, TT) coordinated by a pair of hinges. Helical segments moving as quasi-rigid domains are colored in blue and red. Residues that act as flexible hinges are colored in green. Spheres represent C_α_ atoms of G37 and G38 and are colored according to the domain in which they are located. Screw axes passing the hinge regions are shown in grey. A screw axis perpendicular to the helix axis indicates a bending-like motion while a screw axis parallel to the helix axis indicates a twisting-like motion. For mixed bending/twisting motions, a larger projection of the screw axis with respect to the helix axis indicates a higher percentage of twisting. **(B)** Probability of all 6 types of hinge bending and twisting motions in POPC and **(C)** TFE/H_2_O. **(D, E)** Probability of each residue as hinge site in the single-hinge (B+T) and double-hinge motions (BB+BT+TB+TT) for peptides in POPC **(D)** and **(E)** TFE/H_2_O.

Due to their reduced H-bond stabilities, combined with extensive shifting between α- and 3_10_-H-bonding (**Figure S4**), and the absence of steric constraints, the V_36_GGV_39_ sites in the C99 TMD are optimally suited to act as hinges, an observation discussed already previously (40, 42, 44, 53, 73). Interestingly, consistent with previous results (53), a second hinge located in the TMD-C, upstream of the ε-sites appearing around residues T_43_VI_45_ was revealed. When acting in combination (motion types BB, BT, TB, TT), both hinges coordinate bending/twisting of the flanks (domain B, i.e. residues I47-M51, and domain A, i.e. residues I31-M35, respectively) with respect to the middle part of the helix (residues V36-V46). The impact of the G38 mutations on local H-bond flexibility (**Figure S4**) also alters the location of the flexible joints coordinating the motions of the flanking segments (**Figures 5D** and **E**). In POPC, hinge propensities clearly shifted from G38 in the WT to G37 for G38P for all types of motions. The shift of a hinge site by one residue correlates with the counter-clockwise reorientation of the ε-sites as also documented in **Figure 4C** by a shift of the swivel angle distribution towards more positive values (also see **Table S1**). The most severe impact on single-hinge location was noticed for the G38L mutant in POPC. Restricted rotational freedom around the L38 in the tightly packed lipid environment eliminates preference for single-hinge bending and twisting around the G_37_G_38_ hinge and enhances bending around the second T_43_VI_45_ hinge. Most importantly, although WT and G38L peptides sample similar regions in the bend/swivel space of ε-site orientations (**Figure 4C**), the backbone conformations contributing to the orientation variability are different. In TFE/H_2_O (**Figure 5E**), anisotropic bending over the G_37_G_38_ hinge was confirmed by the equal hinge propensity of these two residues in the WT peptide, while both mutants slightly preferred G37.

In conclusion, the simulations show that the heterogeneous distribution of flexibility in the C99 TMD provides several hinge regions coordinating bending and twisting motions. The helix deformations associated with these motions and the location of the hinges are determined by sequence (WT vs. G38L or G38P) as well as by packing constraints imposed by the environment (POPC membrane vs. TFE/H_2_O). These hinge motions are favored by the absence of steric constraints as well as by flexible H-bonds shifting between i,i+3 and i,i+4 partners. Thus, although the ε-sites reside in a stable helical domain, they possess mobility due to a variety of backbone motions around two flexible regions acting as hinges, the V_36_GGV_39_ region and the T_43_VI_45_ region.

### G38 mutants do not alter contact probabilities with γ-secretase

Since the G38 mutations are localized close to potential TMD-TMD interaction interfaces (i.e. G_29_XXXG_33_, G_33_XXXG_37_ and G_38_XXXA_42_ motifs (40, 43)), an alternative rationale for the impaired γ-secretase cleavage of the G38 mutants (**Figure 1**) could be altered contact preferences with γ-secretase. To screen for contact interfaces of the C99 TMD with γ-secretase, we set up an *in silico* docking assay for transmembrane components (DAFT, (85)) using a coarse-grained description of POPC lipids, water, C99 TMD (C99_26-55_) and γ-secretase (Supporting Information). This protocol was previously shown to reliably reproduce experimentally verified protein-protein interactions in a membrane (85, 86). The use of >750 replicas per TMD starting from unbiased non-interacting initial states, sampling for at least 1 μs per replicate and inclusion of low-amplitude backbone dynamics of γ-secretase (87) provided exhaustive sampling of potential contact sites and exceeds previous assessments of C99 binding sites with respect to both the number of replicates and simulation time (88, 89).

Our calculations show that the C99_26-55_ WT and both mutant peptides could principally interact with the surface of the γ-secretase complex and contact all four complex components (**Figure 6A**). Interestingly, in agreement with previous substrate-crosslinking experiments (21), the C99_26-55_ showed the highest binding preference for the PS1 NTF (**Figure 6A**). The normalized C99_26-55_ proximities for each residue of γ-secretase (**Figure 6B**) revealed that contacts with the highest probabilities were formed between the juxtamembrane S26NK28 residues of C99_26-55_ and the two threonine residues T119 and T124 in the hydrophilic loop 1 between TMD1 and TMD2 of PS1 (**Figure S7**). This observation indicates that the presenilin TMD2 may represent a major exosite of γ-secretase. Interactions between PS1 TMD2 and the C99 TMD could mainly be attributed to contacts of the A30, G33, G37 and V40 of the C99 TMD with residues L130, L134, A137, and S141 of PS1 TMD2 (**Figure 6C** and **Figure S7**). Additional contact sites of the C99 TMD at V44 and I47 are located on the same face of the C99 TMD helix as the main contact sites (**Figure 6C**). However, the probabilities of the observed dominant contacts with the enzyme were not significantly altered by the G38 mutations. Thus, based on our substrate docking simulations, the structural alterations of the C99 G38 mutant TMD helices may not cause gross alterations in initial substrate-enzyme interactions.

**Figure 6.**
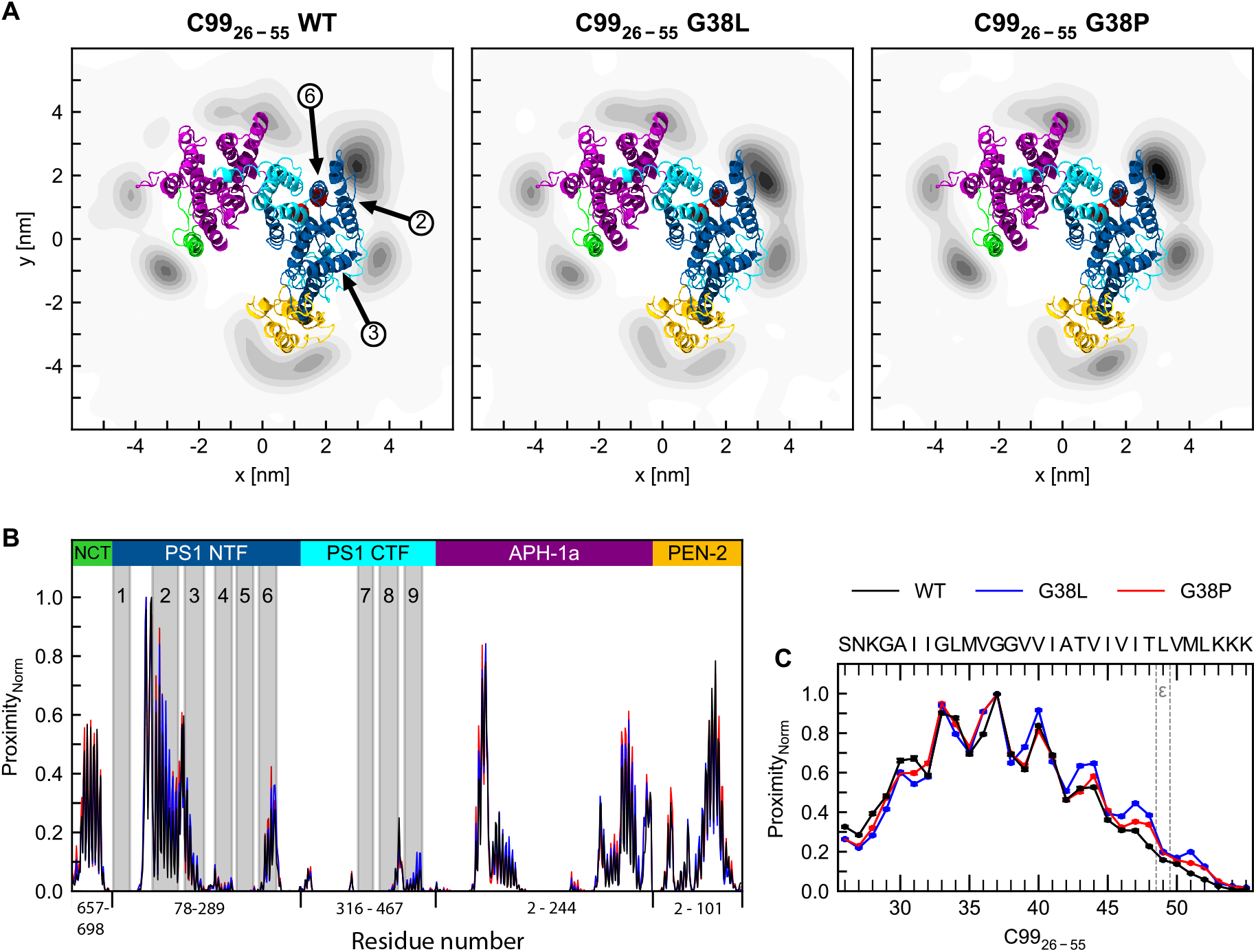
Probability of initial contacts of C99_26-55_ peptides with γ-secretase is not altered for G38 mutants compared to WT in in silico modeling of the encounter complex. **(A)** Kernel densities of the center-of-mass location of the C99_26-55_ peptide. Darker colors indicate higher contact probabilities. The representation shows the parts of γ-secretase that are embedded in the membrane, pertaining to the subunits NCT (green), PS1 NTF (blue), PS1 CTF (cyan) APH-1a (purple) and PEN-2 (yellow). Black arrows highlight TMD2, TMD3 and TMD6 of PS1, the active site aspartate residues in PS1 TMD6 and TMD7 are indicated by red spheres. **(B)** Normalized proximities between residues of γ-secretase subunits and the C99_26-55_ peptide. Grey areas indicate residues that are part of the indicated TMDs of PS1. **(C)** Normalized proximities between C99_26-55_ residues and TMD2 of PS1.

## DISCUSSION

Since conformational flexibility of the substrate may play a key role for substrate recruitment and cleavage by γ-secretase, we focused in our study on the influence of altered intrinsic structural and dynamical properties of the C99 TMD on its cleavability by γ-secretase. In particular, we asked whether cleavability could be correlated with the structural and dynamical properties of the TMD and, if so, what kind of properties would be functionally relevant. As indicated by previously determined NMR structures in detergent micelles (73, 90), as well as by MD simulations in membrane bilayers (40, 42, 43, 91, 92), the C99 TMD contains a flexible hinge region at the G_37_G_38_ residues. Since substrate entry of C99 into the active site of γ-secretase may involve swinging-in of the TMD-C containing the cleavage region (45), we hypothesized that the bending flexibility induced by the G_37_G_38_ hinge could possibly play an important role for the cleavage of this γ-secretase substrate (41, 44). To investigate this issue further, artificial mutations were investigated, designed with the aim to severely alter TMD flexibility by either stabilizing or destabilizing it. CD and solution NMR experiments as well as MD simulations confirmed our rationale that exchanging G38 with leucine leads to a more stable helix, while the G38P mutation reduced helicity. Strongly reduced cleavage efficiency was observed for both mutants in the C99 γ-secretase cleavage assay. This was remarkable since we previously observed that cleavage of C99 was not impaired and rather enhanced when G38 is replaced by the photocrosslinkable, unnatural and bulky amino acid *p*-benzoyl-L-phenylalanine, suggesting that, in principle, this position within the C99 TMD is quite tolerant to structural modifications (21). Interestingly, the processivity of γ-secretase was also dramatically altered for both G38 mutants in a distinct way. These observations show that the G38 mutations have a dramatic impact on both the initial cleavability at the ε-site and the subsequent carboxy-terminal trimming by γ-secretase.

A possibility to explain the impaired cleavage of both G38 mutants of the C99 substrate would be an altered substrate encounter. The ssNMR measurements and corresponding MD simulations revealed a moderate tilt angle and no significant differences between the G38 mutants and WT with regard to the predominant tilt angle of the C99_26-55_ peptide in a bilayer. Additionally MD simulations of the initial contact sites between the C99_26-55_ peptide and γ-secretase in a POPC bilayer did not disclose major differences between the G38 mutants and WT. Interestingly, consistent with previous studies (21), particularly the PS1 NTF subunit of γ-secretase was found as major contact region. However, substrate contacts in the catalytic cleft of the γ-secretase complex were not found. It is likely that relaxations of the enzyme-substrate complex after binding as well as the substrate transfer to the active site take more time than the ~370 μs total simulation time per peptide used in this analysis.

Alternatively, we assessed whether alterations in the backbone dynamics in the proximity of the residues at the ε-sites, caused by the G38 mutations, would give insight on the altered cleavability of the mutants. DHX and HDX experiments that report on the stability of H-bonds in the C99 TMD revealed differences between WT, G38L, and G38P, in agreement with the observed structural changes. Thus, compared to WT, the overall D/H exchange kinetics was slower for G38L and faster for G38P, as expected. A more detailed residue-specific analysis revealed that the effect of the G38 mutations on D/H exchange only occurred at the residues in the vicinity of the G_37_G_38_ hinge while residues in the proximity of the ε-sites were not affected. For these sites, MS-ETD-DHX, NMR-HDX and MD simulations consistently reported low exchange rates. MD simulations further revealed protection of the backbone amides at the ε-sites by stable intrahelical H-bonds. Mutation-induced loosening of the H-bonds or shifting populations of α- and 3_10_-H-bonds was not observed at these sites. These observations show that backbone dynamics at the ε-sites are not affected by the G38 mutations and can thus not explain the reduced cleavage by γ-secretase of both the G38L and the G38P mutation of C99. Interestingly, D/H exchange rates at the γ-cleavage sites of the C99_26-55_ peptide were decreased for the G38L mutation and slightly increased for G38P compared to WT. However, these findings cannot be applied to explain the altered processivity of the mutants, since the backbone dynamics of the shortened C99 TMD may change after the AICD has been cleaved off.

In general, α-helical structures of TMDs are stabilized by van der Waals interactions, as well as a strong H-bond network between i,i+3 and i,i+4 neighbors. Remarkably, for the C99 TMD, water accessibility in the TMD-C has been reported to be extremely limited (40), consistent with our findings of a very high degree of H-bond occupancy and extremely slow D/H exchange in the TMD-C found here in this study as well as previously for other C99 TMD derived segments (43, 52). Interestingly, in the TFE/H_2_O environment amide exchange is nevertheless extremely slow indicating that the helix around the ε-sites is quite stable. Thus, helix instability is not an intrinsic property of the TMD-C at all, and is also not modulated by our G38 mutations. It appears that opening of the α-helix of the residues in the vicinity of the ε-sites might be the major hurdle for substrate cleavage and is induced only upon an interaction with the enzyme. Although these results confirm previous studies (43, 44, 52), they are in contrast with a recent study by Cao et al (93), who calculated D/H fractionation factors from ratios of exchange rates in order to determine H-bond strengths of C99 in lysomyristoyl-phosphatidyl glycerol micelles. However, as explained elsewhere (68), H-bond strengths derived by the approach of Cao et al (93) describe the preference for deuterium in an amide-to-solvent H-bond rather than the properties of the intrahelical amide-to-carbonyl H-bonds. Furthermore, in this environment the C99 TMD was reported to be artificially kinked (42, 73, 90).

Models of enzymatic substrate processing provide evidence that conformational dynamics of substrates and enzymes play a key role for recognition and relaxation steps (94-97). Here, the intrinsic dynamics encompass all the conformations necessary for substrate binding, pre-organization of the substrate-enzyme complex and the stabilization of the transition state. The chemical reaction is thought to be a rare, yet rapid, event (98), that occurs only after sufficient conformational sampling of the enzyme-substrate complex to generate a configuration that is conducive to the chemical reaction. For the γ-secretase/substrate complex this sampling could be at the level of substrate transfer from exosites to the active site as well as at the level of substrate fitting into the active site. A series of relaxations and mutual adaptation steps of substrate and enzyme after initial binding might be required before the scissile bond fits into the active site. Thus, multiple conformational selection steps may play a decisive role in distinguishing between substrates and non-substrates (96). Here, large-scale shape fluctuations might be selected to enable recognition, while lower-amplitude, more localized motions help to optimize and stabilize the enzyme-bound intermediate states (94-97, 99). With regard to C99, the essential property enabling the TMD to switch between different shapes of functional importance is the organization of rigidity/flexibility along the helix backbone where residues enjoying higher flexibility can coordinate the motions of more rigid flanking segments. These flexible hinges might provide the necessary bending and twisting flexibility for orienting the reaction partners properly. Our MD simulations here and previously reveal that the residues T43VI45 upstream of the ε-sites provide additional hinge flexibility which may be of importance for conformational adaptation of the TMD to interactions with the enzyme where large-scale bending is obstructed (53). In addition to bending around the G3_7_G_38_ sites, twisting and more complex collective motions including combinations of bending and twisting around the pair of hinges can occur. This structural distribution translates into a diversity of ε-site orientations. The perturbation of this distribution might provide plausible explanations of the reduced cleavability of both, the G38L and the G38P mutant. Particularly, the counterclockwise shifts of the orientation of the ε-sites for both G38L and G38P mutants in POPC and the absence of small bending angles for G38P indicate that presentation of the scissile bond to the active site of presenilin can be misdirected for each mutant in its own way and differently compared to WT. Thus, the modified mechanical linkage of helical segments constituting the substrate TMD might increase the probability of non-productive interactions with the enzyme what could explain the reduced cleavability of the G38 mutants. A similar mechanism was proposed recently to explain reduced ε-cleavage efficiency induced by FAD mutations upstream of the ε-sites in the C99 TMD (53, 68).

Notably, the TMD of Notch1, as well as the TMD of the insulin receptor (IR), two other substrates of γ-secretase (9), have conformations very different from that of the C99 TMD as determined from solution-NMR of single-span TM helices in membrane mimics (100, 101). In particular, Notch1 appears to be a straight helix, while the TMD helix of the insulin receptor is S-shaped resembling the minor population of double-hinge conformations of the C99 TMD in a POPC bilayer found in our study. These observations seem to challenge the swinging-in model coordinated by a central hinge as an integral step for the passage of the substrate toward the active site. Nevertheless, generally the conformational repertoire of the TMD of the substrates is determined by α-helical folding where helices bend and twist around several sites. The relative importance of the individual conformations reflects differences in local flexibility. Functionally relevant conformations are not necessarily among the highest populated ones. Rather conformations for which the protein has a low intrinsic propensity might be selected for productive interactions with the enzyme regulating or coordinating mechanistic stages preceding catalysis. These so-called ‘hidden’, ‘invisible’ or ‘dark’ states are amenable by NMR or MD methods (99, 102-104). Although missing large-scale helix bending, Notch1 and IR (and even other substrates of γ-secretase) might nevertheless provide the repertoire of functionally important motions necessary to adapt to interactions with the enzyme at different stages of the catalytic cycle. Furthermore, binding and conformational relaxation steps of different substrates might follow different pathways to optimize the catalytic competent state (95, 96).

## CONCLUSION

Taken together, since this study reveals that the G38 mutations do not impact the structural and dynamical properties around the ε-cleavage site but nevertheless do have a severe impact cleavage and processivity of the C99 substrate by γ-secretase, we conclude that necessary conformational relaxations required to facilitate the proteolytic event at the active site are not due to intrinsically enhanced flexibility of the C99 substrate around its ε-cleavage site but must be induced by interactions of the substrate with the enzyme. Interestingly, in line with this interpretation of our data, it was recently concluded, based on vibrational spectroscopy and NMR studies of enzyme-substrate interactions, that PSH, an archaeal homolog of presenilin, can induce local helical unwinding towards an extended β-strand geometry in the center of the TMD of the substrate Gurken (105) as well as in the ε-cleavage site region of a C99 TMD derived substrate (106). Nevertheless, our study suggests that, prior to the catalytic event, intrinsic conformational flexibility of substrate and enzyme is also necessary to prepare access to the cleavage site and orient the reaction partners properly. As conformational adaptability of the C99 substrate TMD is provided by flexible regions coordinating motions of helical segments, subtle changes of H-bond flexibility induced around the G_3_7G_38_ hinge by G38 mutations alter the local mechanical linkage to other parts of the helix. As a consequence, irrespective of whether the mutation at the G_37_G_38_ hinge is helix stabilizing or helix destabilizing, the orientation of remote initial cleavage sites can be misdirected in such a way that the probability of productive orientations with the active site of γ-secretase is decreased, leading to impaired cleavage and altered processivity.

## AUTHOR CONTRIBUTIONS

D.H., B.L., D.L., C.S., C.M.-G., H.S. conceived the study, designed experiments, analysed and interpreted data, and supervised research. A.G. and S.M. performed MD simulations, F.K. CD spectroscopy, N.M. cleavage assays, P.H. hydrogen/deuterium exchange experiments, and M.S., H.H., A.V., H.F. NMR spectroscopy. A.G., S.M., F.K., N.M., P.H., M.S., H.H., A.V., H.F. analysed and interpreted data. F.K. coordinated the drafting of the manuscript. F.K., C.M.-G., A.G., C.S. and H.S. wrote the manuscript with contributions of all authors.

## ACKNOWLEDGEMENTS

This work was supported by the DFG (FOR2290) (D.H., B.L., D.L., C.S., H.S.) and in part by the VERUM foundation (F.K.). Computing resources were provided by the Leibniz Computing Center through grant p292so as well as by the Gauss Centre for Super Computing through grant pr48ko. We thank Martin Zacharias for providing results of allatom simulations of γ-secretase and Marius Lemberg for critical reading of the manuscript

